# Deciphering the genetic basis of wheat seminal root anatomy uncovers ancestral axial conductance alleles

**DOI:** 10.1101/2020.11.19.389882

**Authors:** Elisha Hendel, Harel Bacher, Adi Oksenberg, Harkamal Walia, Nimrod Schwartz, Zvi Peleg

## Abstract

Root axial conductance which describes the ability of water to pass throw the xylem, contributes to the rate of water uptake from the soil throughout the whole plant lifecycle. In a rainfed wheat agro-system, grain-filling is typically occurring during declining water availability (i.e. terminal drought). Therefore, preserving soil water moisture during grain filling could serve as a key adaptive trait. We hypothesized that lower wheat root axial conductance can promote higher yields under terminal drought. A segregating population derived from a cross between durum wheat and its direct progenitor wild emmer wheat was used to underpin the genetic basis of seminal root architectural and functional traits. We detected 75 QTL associated with seminal roots morphological, anatomical, and physiological traits, with several hotspots harboring co-localized QTL. We further validated the axial conductance and central metaxylem QTL using wild introgression lines. Field-based characterization of genotypes with contrasting axial conductance suggested the contribution of low axial conductance as a mechanism for water conservation during grain filling and consequent increase in grain size and yield. Our findings underscore the potential of introducing wild alleles to reshape the wheat root system architecture for greater adaptability under changing climate.

## Introduction

Plant roots are the route for water and nutrients acquisition from the soil, and provide physical anchoring, photoassimilates storage, and interface with the rhizosphere. The root system architecture is responsive to spatiotemporal soil factors such as moisture and minerals availability, temperature and, pH (Robbins & Dinneny, 2015). Root system characteristics including morphological and anatomical structure, carbon allocation plasticity, hydro- and gravitropism, and the rhizosheath, play a crucial role in the plant’s adaption and response to various environmental cues (Kitomi et al., 2020; Klein, Schneider, Perkins, Brown, & Lynch, 2020; Lynch, 2018; Rellan-Alvarez, Lobet, & Dinneny, 2016; Tracy et al., 2020; Uga et al., 2013). A better understanding of the root system genetic architecture can facilitate the development of new cultivars that ameliorate water-use efficiency for the projected climate change (Gupta, Rico-Medina, & Caño-Delgado, 2020; Palta & Turner, 2019; Preece & Peñuelas, 2020; Schneider & Lynch, 2020).

Water absorption from the soil to the transpiring leaf is mediated via complex anatomical-structural pathways, starting with the epidermis layer and the root hairs (tubular outgrowths of epidermal cells). Root hairs extend the absorbing surface by more than 50% and play an essential role in nutrient acquisition and water uptake from the soil (Carminati et al., 2017; Dolan, 2017). After entering the root, water moves via either the apoplastic (within parenchyma cell walls) or symplastic route (crossing through cell membranes) towards the root center. When the water reaches the endodermis (outer layer) of the root cylinder, it encounters the casparian bands, which are suberin cell walls that force the water to move via the symplastic route until it is loaded into the xylem (Holbrook, 2018). Xylem is made up of hollow and dead cells-tracheae, which are positioned one on top of each other with a perforation plate between them, creating a long tube structure that continues through shoot and leaves.

Wheat (*Triticum* sp.) is a staple grain-crop, with an annual production of ∼760 million tons worldwide. Wheat domestication and subsequent evolution under domestication involved a suite of complex genetic, morphological, anatomical, and physiological modifications (Abbo et al., 2014; Golan, Hendel, Méndez Espitia, Schwartz, & Peleg, 2018). Wild emmer wheat [*T. turgidum* ssp. *dicoccoides* (Körn.) Thell.], the direct allo-tetraploid (2n=4x=28; genome BBAA) progenitor of domesticated wheats, thrives across a wide range of eco-geographic habitats through the Fertile Crescent and harbors an ample allelic repertoire for agronomically important traits, including drought tolerance (e.g. Bacher et al., 2020; Golan et al., 2018; Peleg et al., 2005; Peleg et al., 2009). During the transition from natural habitat to anthropogenic agro-systems, ∼10,500 years ago, many adaptive traits were lost or gradually eliminated from the domesticated germplasm.

The monocotyledonous wheat root system is composed of two types of roots: seminal roots (i.e. embryonic), and adventitious roots (i.e. crown or nodal) (Klepper, Belford, & Rickman, 1984). The seminal root system develops initially and consists of the primary (or radicle) root and two pairs of seminal roots (#2-3 and #4-5) that develop at the scutellar node of the embryonic hypocotyl (Percival, 1921). Seminal roots penetrate the soil earlier and deeper than the adventitious roots and usually remain active throughout the whole plant life cycle, and play a crucial role in absorbing water from deep soil layers (Locke & Clark, 1924; Watt, Magee, & McCully, 2008; Weaver, 1926). Seminal root xylem in wheat is usually patterned with one large metaxylem element in the center (CMX), and peripheral elements surrounding it in one or two rings. As the root develops, some of the peripheral elements can increase in size becoming similar to the initial CMX. In recent years, several studies on root system architecture uncovered the genetic basis of these traits, however, most studies focused on the geometric traits, such as root number, length, diameter and, angle (Christopher et al., 2013; Golan et al., 2018; Hamada et al., 2012; Shorinola et al., 2019; Soriano & Alvaro, 2019; Voss-Fels, Snowdon, & Hickey, 2018). On the other hand, root anatomical traits were less studied and remain largely unexplored.

Under the semi-arid Mediterranean climate, grain maturation period usually occurs under terminal drought. Therefore conserving a larger proportion of soil water moisture during the vegetative phase, is critical to support the subsequent grain filling. It has been suggested that restricting root hydraulic conductance can induce stomatal closure and minimize water loss (Vadez, Kholova, Medina, Kakkera, & Anderberg, 2014). In accordance, Richards and Passioura (1989) hypothesized that reducing wheat seminal roots xylem element size can decrease the axial conductance (Kx) and thereby decelerating early water-use. Here, we present evidence that lower Kx supports sustainable grain yield under terminal drought. Further, we targeted Kx-related morphological and anatomical seminal root traits and uncovered their underlying genetic basis, using recombinant-inbred lines population, derived from a cross between durum wheat and wild emmer. Lastly, we show that introgression of major genomic regions for Kx and central metaxylem diameter in modern durum wheat background can reshape its root hydraulic properties. The identified genomic regions may serve as a basis for future breeding efforts in developing wheat cultivars adapted to the projected climate change.

## MATERIALS AND METHODS

### Plant material

A mapping population of 150 F_8_ recombinant inbred lines (RILs) was developed by single-seed descent from a cross between an elite durum wheat (♀) cultivar Svevo and the wild emmer wheat (♂) accession Zavitan, as was previously described (Avni et al., 2014). For QTL validation we used a set of introgression lines (ILs), derived from adaptive RILs [i.e. genetic composition of post-domestication alleles: reduce height (*Rht-B1*) and non-brittle spike (*Tdbtr*), that were backcrossed three times with Svevo follow by five generations of self, to generate BC_3_F_5_].

### Seedlings growth conditions

Seedlings were grown using the ‘cigar roll’ method (Watt et al., 2013). Twelve uniform-size seeds of each line were placed on a moist germination paper (25 × 38 cm; Anchor Paper Co., St. Paul, MN, USA), about 2 cm apart, with the germ end facing down. The paper was covered with another sheet of moist germination paper rolled to a final diameter of 3 cm. The bases of the rolls were placed on a tray, with half-strength Hoagland solution, in a darkened growth chamber at a constant temperature of 16.5°C. The nutrient solution contained the following macro elements: KNO_3_ (3mM), Ca (NO_3_)_2_ (2mM), KH_2_PO_4_ (1mM), MgSO_4_ (0.4mM), and micronutrients: KCl (50µM), EDFS (30µM), H_3_BO_3_ (25µM), ZnSO_4_ (2µM), MnSO_4_ (2µM), H_2_MoO_4_ (0.75µM), CuSO_4_ (0.5µM), pH=6.0. The root hair experiment was conducted after four days in the darkened growth chamber. The anatomical cuts and root length experiments were moved to a daily cycle of 10/14 hours of light and darkness at 16 and 24°C respectively, for three more days, in total 7 days after sowing. During the additional three days, the light intensity was 150-200 (µmol s^-1^ m^-1^).

### Characterization of seminal roots anatomical and physiological traits

Anatomical parameters of all experiments were taken using freehand sections as described previously (Zelko et al., 2012). The primary roots of 7-days old seedlings from each genotype (*n*=4) were cut at the root tip (1.5 cm above the tip) and root base (1.5 cm under the root-to-shoot junction) (Fig. 1). Each segment was embedded in agarose (5%) that was heated beforehand. After the agarose was hardened root sections were cut vertically using a razor, and stained with Toluidine blue (0.0025% for 2 min). Sections were visualized using a Zeiss Axioplan equipped with an Axiocam 105 color camera (Zeiss, Germany). *Root diameter* (the diameter of the root circle), c*ylinder diameter* (the diameter of the root stele), *central metaxylem area* (CMX), and *peripheral xylem area* (PX, consisting of protoxylem and other xylem elements except for CMX) (see Fig. 1) were measured using Image J software (https://imagej.net/ImageJ_Ops). The transformation from diameter to area and *vice versa* were based on the assumption of perfectly cylindrical elements (Tyree & Ewers, 1991), for example in the *CMX area to CMX diameter*.

**Figure 1.**
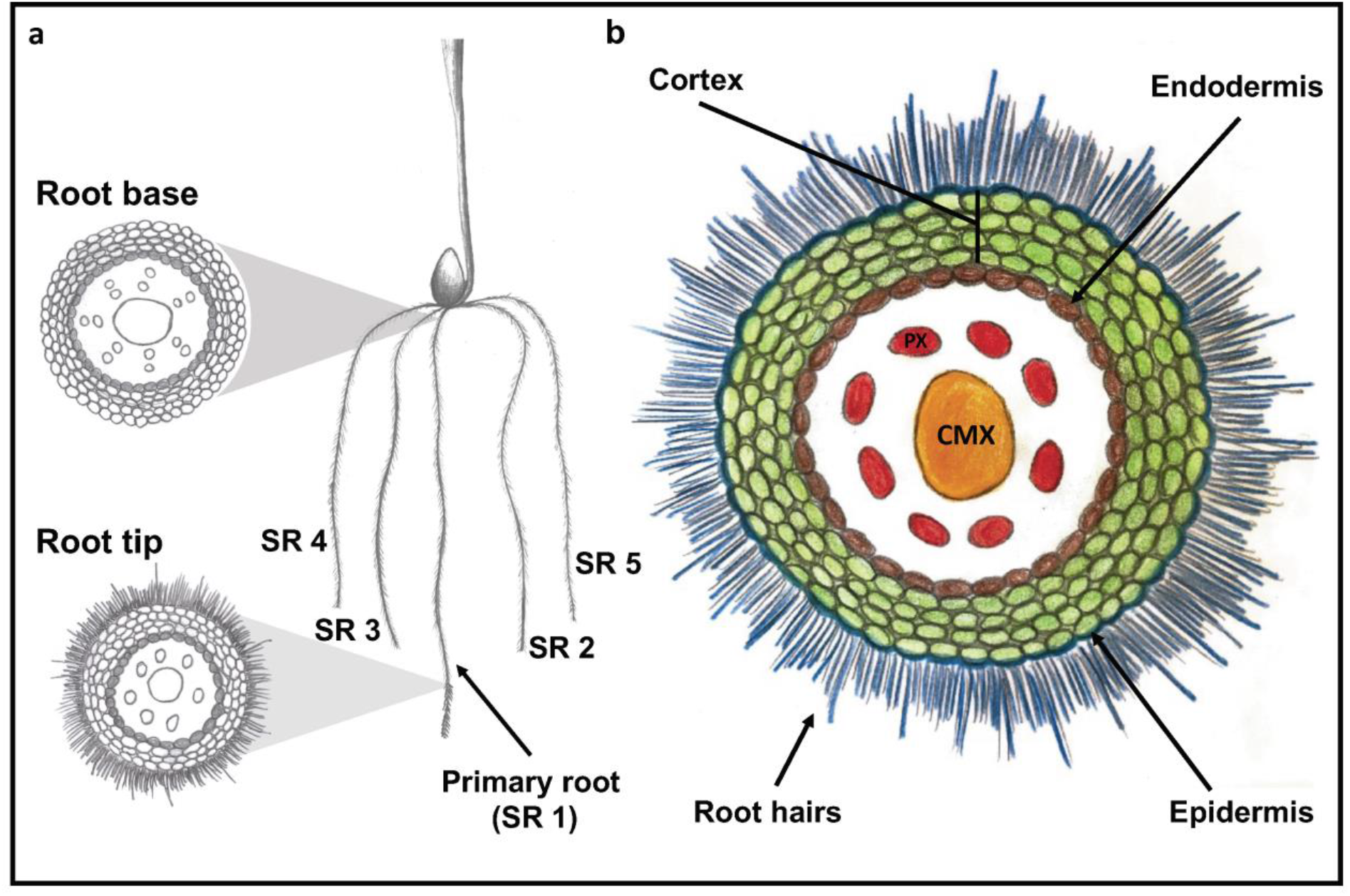
A schematic illustration of wheat seminal root apparatus and anatomical cross-section. (**a**) The primary seminal root (SR1), the two symmetric root pairs (SR2,3 and SR4,5), and the location of root base and root tip cross-sections. (**b**) Root tip anatomy consists of central metaxylem (CMX), peripheral xylem (PX), endodermis, epidermis, cortical parenchyma (Cortex), and root hairs.

Based on the measured traits we calculated the *total xylem area, cylinder to whole root ratio, CMX to total xylem area ratio, xylem to whole root ratio*, and *xylem to cylinder area ratio* and as presented in the following equations:

1. *Total xylem area = CMX area + PX area*
2. 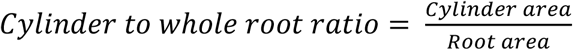
3. 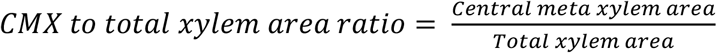
4. 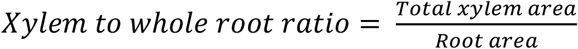
5. 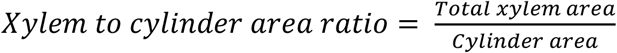

*Axial conductance* (Kx) was calculated using modified Hagen–Poiseuille’s equation for fluid flow through a bundle of the perfectly cylindrical pipe as described previously (Tyree & Ewers, 1991):

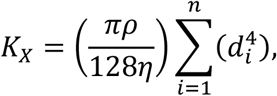

where k_x_ (kg s^-1^ m MPa^-1^) is the hydraulic conductance, *ρ* is the density of the fluid (for distilled water, 1,000 kg·m^−3^), *η* is the dynamic viscosity of the fluid (for distilled water 10^−9^ MPa·s), *d* is the diameter (m) of the *i*^th^ pipe, and *n* is the number of pipes. The calculation of axial conductance based on anatomical root cross-sections results in a theoretical Kx which is a common method (e.g., Melchior & Steudle, 1993; Frensch & Steudle, 1989; Tyree & Zimmermann, 2002; Olson & Rosell, 2013).

### Field-based characterization of Kx, and yield components

Two genotypes with contrasting hydraulic properties: Svevo (high Kx) and IL82 (low Kx) were selected for this assay. Svevo is an elite Italian durum wheat cultivar released in 1996 (CIMMYT line/Zenit) is the reference variety for quality and productivity of durum wheat. IL82 consists 13.45% (15 introgressions) of the genome from wild emmer wheat accession Zavitan on the background of Svevo. A paired sample experimental design between the two genotypes with two irrigation regimes: well-watered and terminal drought stress, was employed, with 5 replicates. Uniform seeds of Svevo and IL82 were sown in an insect-proof screen-house protected by a polyethylene top, during the winter of 2019-2020 at the experimental farm of The Hebrew University of Jerusalem in Rehovot, Israel (34°47′N, 31°54′E; 54 m above sea level). The soil at this location is brown-red degrading sandy loam (Rhodoxeralf) composed of 76% sand, 8% silt, and 16% clay. Each plot consisted of four rows, with 10 plants 10 cm apart where the circumference plants of each plot served as borders, resulting in a total of 16 plants examined for each plot. The field was treated with fungicides and pesticides to avoid the development of fungal pathogens or insect pests, and was weeded manually once a week. Two irrigation regimes were applied via a drip system: well-watered and terminal drought mimicking the natural pattern of rainfall in the Eastern Mediterranean basin; water was applied during the winter months starting from planting (December 12, 2019) and ending on April 23. Well-watered treatment was irrigated twice a week throughout the season. The terminal drought treatment started at the phenological stage 2 to 3 tillers [Zadoks growth scale 22-23 (Zadoks, Chang & Konzak, 1974)] and was irrigated twice every other week. The seasonal water-use, including irrigation and soil water depletion, was ∼750 and ∼250 mm for the well-watered and terminal drought treatments, respectively.

Anatomical traits from field experiment have been taken with additional two steps: *i*) the top of the root system has been harvested with a small shovel]i.e. “mini-shovelomics” (Trachsel, Kaeppler, Brown, & Lynch, 2011)], and *ii*) adventitious roots had been removed manually to expose the seminal roots (Fig. S1). From this stage, the protocol for calculating anatomical traits was the same as above. At the end of the experiment, each plot was harvested, spikes were separated from the vegetative organs (stems and leaves) and both plant parts were oven-dried (80°C for 48 h) and weighed. Spikes were then threshed with laboratory seed thresher (Wintersteiger, LD-350), weighed and counted to obtain grain yield (GY) and thousand kernel weight. Relative grain yield was estimated for each genotype as the ratio between each plot under terminal drought and the average performance under well-watered.

### Morphological characterization of seminal roots traits

Whole root systems of seven-days old seedlings (Zadoks stage 10) were scanned in a flatbed scanner (Avision, FB5000), and *primary root length* was measured (*n*=3) using the RootNav software (Pound et al., 2013). To analyze the root hair related traits, the primary root of a 4-days old seedling (Zadoks stage 09) was imaged above the elongation zone using a stereomicroscope (SZX16, Olympus, Tokyo, Japan) equipped with a DP73 digital camera. *Root hair length* was calculated as the average of 10 individual hairs in each photo using ImageJ software (*n*=3) (Fig. S2a). *Root hair density* was analyzed based on an index (range 1-6; Fig. S2b) and scored manually by a panel of nine peoples.

### QTL mapping correspondence hotspots and genomic dissection

A genetic linkage map of 2,110 cM long with an average distance of 0.92 cM was previously developed using the 90K iSelect SNP assay (Illumina) as described (Avni et al., 2014). The QTL analysis was performed with the MultiQTL software)ver. 4.6) using the general interval mapping for a RIL-selfing population, as previously described (Peleg et al., 2009). In brief, the entire genome was screened for genetic linkage, using single-trait analysis, then multiple interval mapping, which incorporates into the model interfering effects of other QTL on a separate chromosome to reduce the residual variation. The hypothesis that a single locus (H_1_) or two linked loci (H_2_) on the considered chromosome have an effect on a given trait was compared against the null hypothesis (H_0_) that the locus had no effect on that trait. Once the genetic model was chosen, 10,000 bootstrap samples were run to estimate the main parameters: locus effect, its chromosomal position, its logarithm of odds (LOD) score and the percentage of explained variance (PEV). For visualization, we used the QTL circos plot created with Circa (http://omgenomics.com/circa).

Correspondence between QTL of different traits was determined using the hypergeometric probability function, as described previously (Peleg et al., 2009):

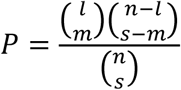

where *n* is the comparable number of intervals, *m* is the number of ‘matches’ (QTL of two traits with >50% overlap of their confidence intervals) declared between QTL, *l* is the total number of QTL found in the larger sample and *s* is the number of QTL found in the smaller sample. Hotspot genomic regions were designated as overlapping of at least three QTL. The genomic region size was considered from the QTL with the smallest standard deviation in the cluster and the other QTL that overlap with its boundaries.

A meta-analysis of overlaps between QTL detected in the current study and previously published information conducted on the GrainGenes genome browser (Blake et al., 2019), where the physical location of the QTL peak was used to locate a gene sequence on the Zavitan genome (Avni et al., 2017), and blast the sequence against Svevo genome (Maccaferri et al., 2019).

### Statistical analyses of phenotypic data and visualization

Statistical analysis was done by JMP^®^ ver. 15 statistical package (SAS Institute, Cary, NC, USA). Descriptive statistics are graphically presented in box plot: median value (horizontal short line), quartile range (25 and 75%), and data range (vertical long line), where t-test was applied between genotypes for all comparisons unless specified otherwise. Dunnett test was used to compare different ILs from their recurrent parent Svevo.

Visualization of phenotypic scaled density and morpho-anatomical phenotypic correlation matrix was done with R Studio (R Core Team, 2021). RStudio: integrated development for R. RStudio. Inc., Boston, MA), using “ggplot” and “corrplot” packages, respectively.

## RESULTS

### Lower axial conductance associates with grain yield under terminal drought conditions

To test the contribution of wild emmer wheat alleles to axial conductance (Kx), we dissected a set of ten wild emmer introgression lines (ILs). In general, most ILs exhibited lower Kx as compared with their recurrent parent Svevo (42.08 *vs*. 50.94 kg s^-1^ m MPa^-1^, Table S1, Fig. S3). We selected IL82 which exhibited significantly lower Kx (33.37 *vs*. 50.94 kg s^-1^ m MPa^-1^; *P*<10^−4^) compared with Svevo (Fig. S3). Characterization of the two genotypes in the field was conducted along with three developmental phases which represent critical stages under the Mediterranean basin climate wheat agro-system. The two first measurements were taken at the beginning of the season, before the application of the water stress. At the seedling stage (Zadoks 11) IL82 exhibited 49% lower Kx compared to Svevo, which was maintained (53%) during the tillering stage (Zadoks 23-25). A similar trend was observed at the end of the season (Zadoks 92) in both treatments, where IL82 showed 36% lower Kx compared to Svevo under well-watered conditions and 31% under terminal drought (Figs. 2 and S4). To address our hypothesis, we compared the grain yield components between Svevo and IL82. Under terminal drought, IL82 had a higher grain yield (*P*=0.059; Table. S2) derived from greater grain size (*P*=0.04; Fig. 2c). In contrast, under well-watered conditions Svevo exhibited a higher grain yield (*P*<0.001). It is worth noting Svevo’s grain yield reduction was ∼two-fold larger compare to IL82, as expresses in relative grain yield (Fig. 2d; Table S2). Image-based phenotyping of Svevo and the IL82 responsiveness to water stress at the vegetative stage showed that both lines had similar biomass and architecture under stress (Bacher et al., 2020).

**Figure 2.**
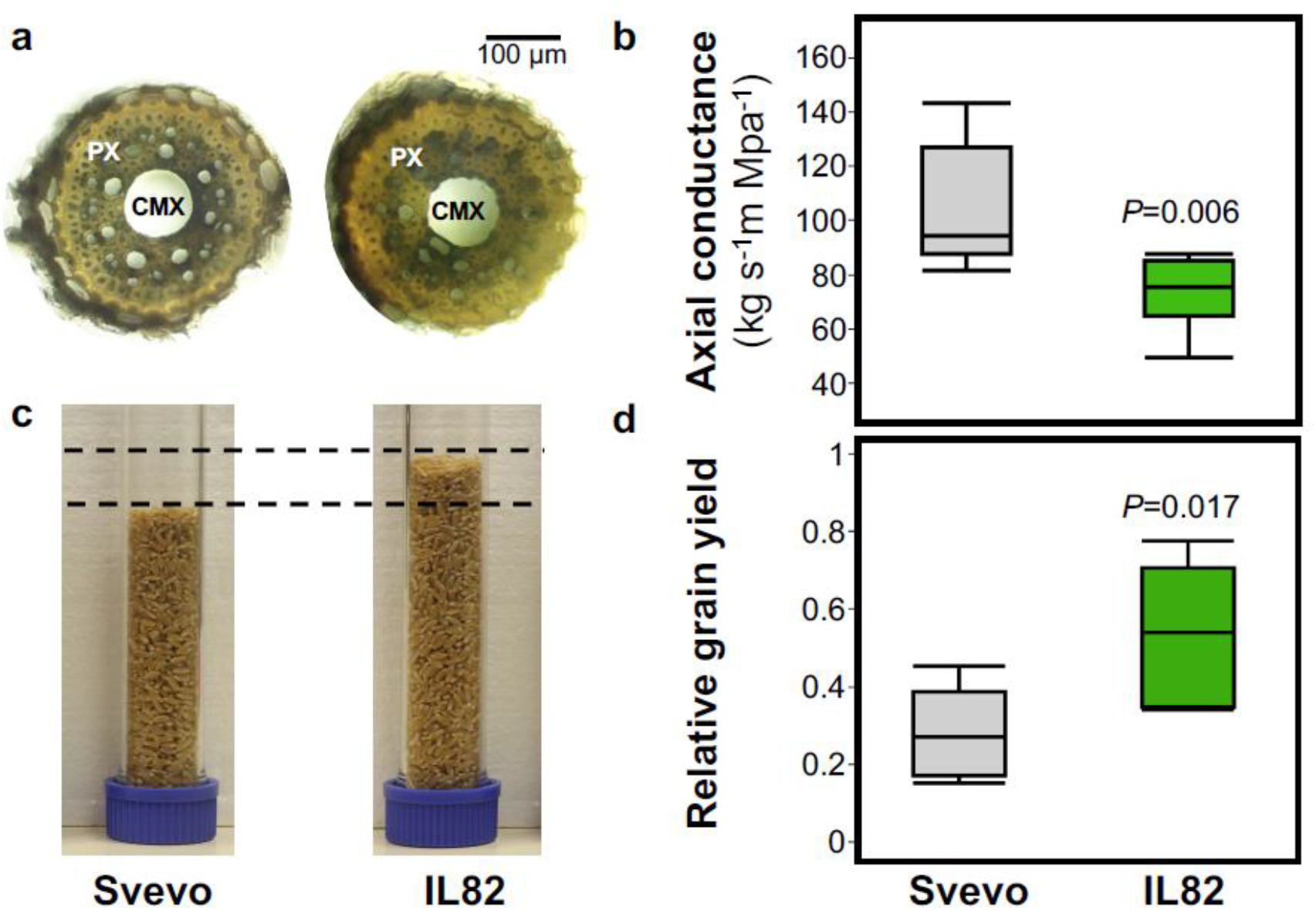
Field-based evaluation of primary seminal root Kx and grain yield under terminal drought conditions. (**a**) Root base anatomical cross-sections. Metaxylem (CMX) and peripheral xylem (PX) elements are indicated. (**b**) Axial conductance of Svevo and IL82 under terminal drought. (**c**) Comparison of two thousand grains between Svevo and IL82 under terminal drought. (**d**) Relative grain yield of Svevo and IL82 under terminal drought compared with well-watered conditions. Differences between genotypes were analyzed by t-test (*n*=5).

### Phenotypic dissection of root morphological, anatomical and physiological traits

The contribution of lower axial conductance (Kx) as a potentially important trait supporting sustainable grain yield production under the Mediterranean climate motivated us to further dissect the genetic and physiological basis of Kx. We used a RIL population derived from a cross between wild emmer and durum wheat to determine the genetic architecture of various seminal root morphological and anatomical traits. Characterization of the population *primary root length* showed strong transgressive segregation (range 134-184 mm), with both parental lines exhibiting similar length (157.94 *vs*. 150.55 mm for Svevo and Zavitan, respectively) (Fig. 3a). Root hairs play important roles in nutrient and water uptake as well as interactions with the rhizosphere by increasing the root volume. In general, the two parents exhibited an opposite trend in *root hair length* and *root hair density*, while Svevo had higher length, Zavitan had higher density and vice versa. Notably, the population showed quantile frequency distribution with most RILs exhibiting similarly to the Zavitan parent (Fig. 3b, c).

**Figure 3.**
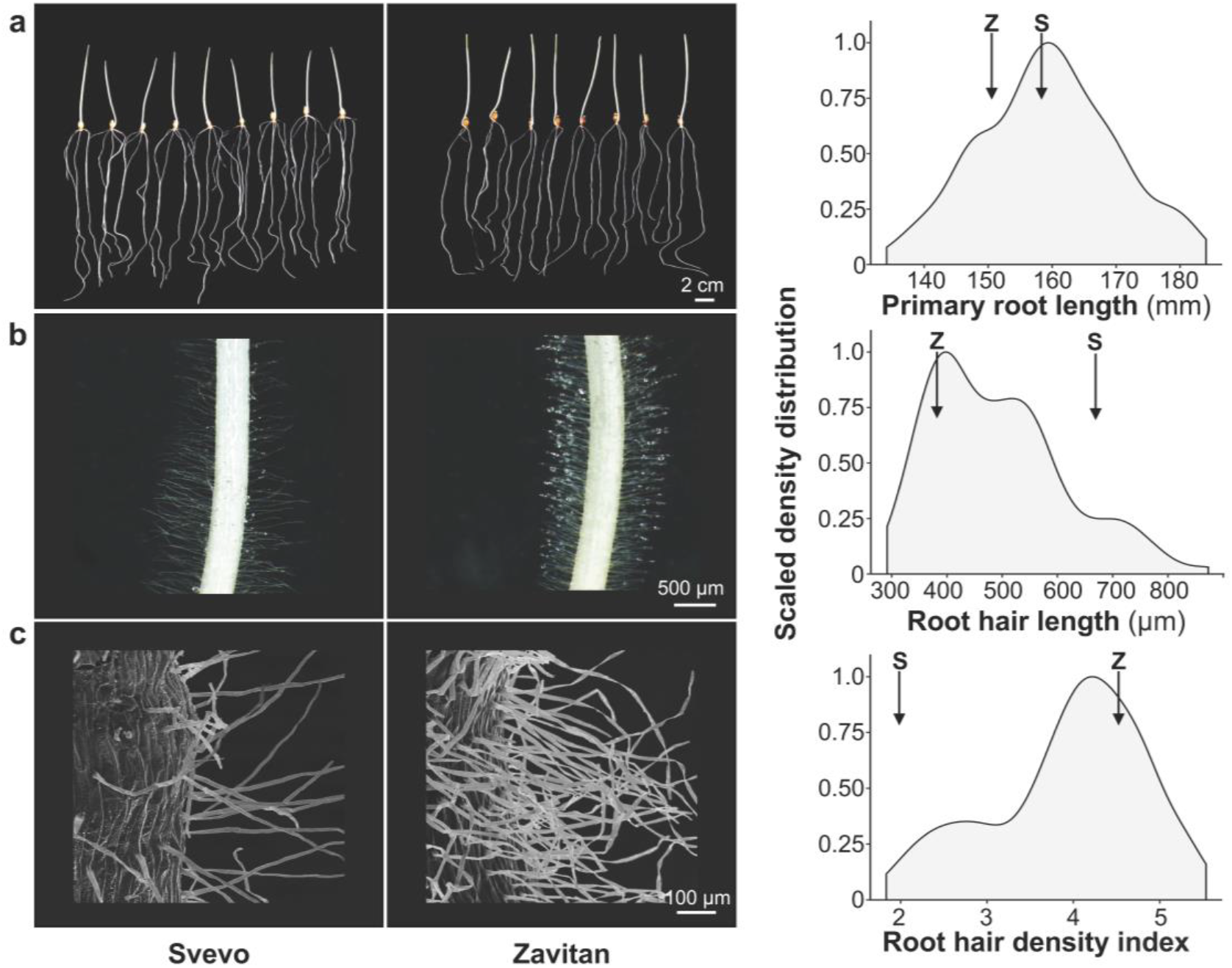
Phenotypic distribution of primary root morphological traits. A representative image of the two parental lines: Svevo and Zavitan, and density distribution of recombinant inbred lines population (Svevo × Zavitan) for (**a**) primary root length, (**b**) root hair length, and (**c**) root hair density index. Values of the parental lines Svevo (S) and Zavitan (Z) indicate by arrows (*n*=3).

To characterize the anatomical structure of the seminal roots, we analyzed two developmental stages of xylem elements: root tip (i.e. middle differentiation process) and base (mature elements). Density distributions of the root tip and base anatomical and physiological traits among the RILs are presented in Figure 4. All variables exhibited normal distribution and most traits showed transgressive segregation. In general, the range of most measured traits (except root diameter) was higher in the root base compared with the root tip, with some level of overlapping (Fig. 4a-e). The two parental lines exhibited different behavior for the root base and tip. While in some cases there was a constant difference between the parents for both root tissues (root diameter, root cylinder diameter, total xylem area, cylinder to whole root ratio, and xylem to cylinder area ratio), in other cases the difference between the parents varied among tissues (PX area and CMX to total xylem area ratio). In two cases the parental lines showed opposite behavior (CMX diameter and Kx) (Fig. 4).

**Figure 4.**
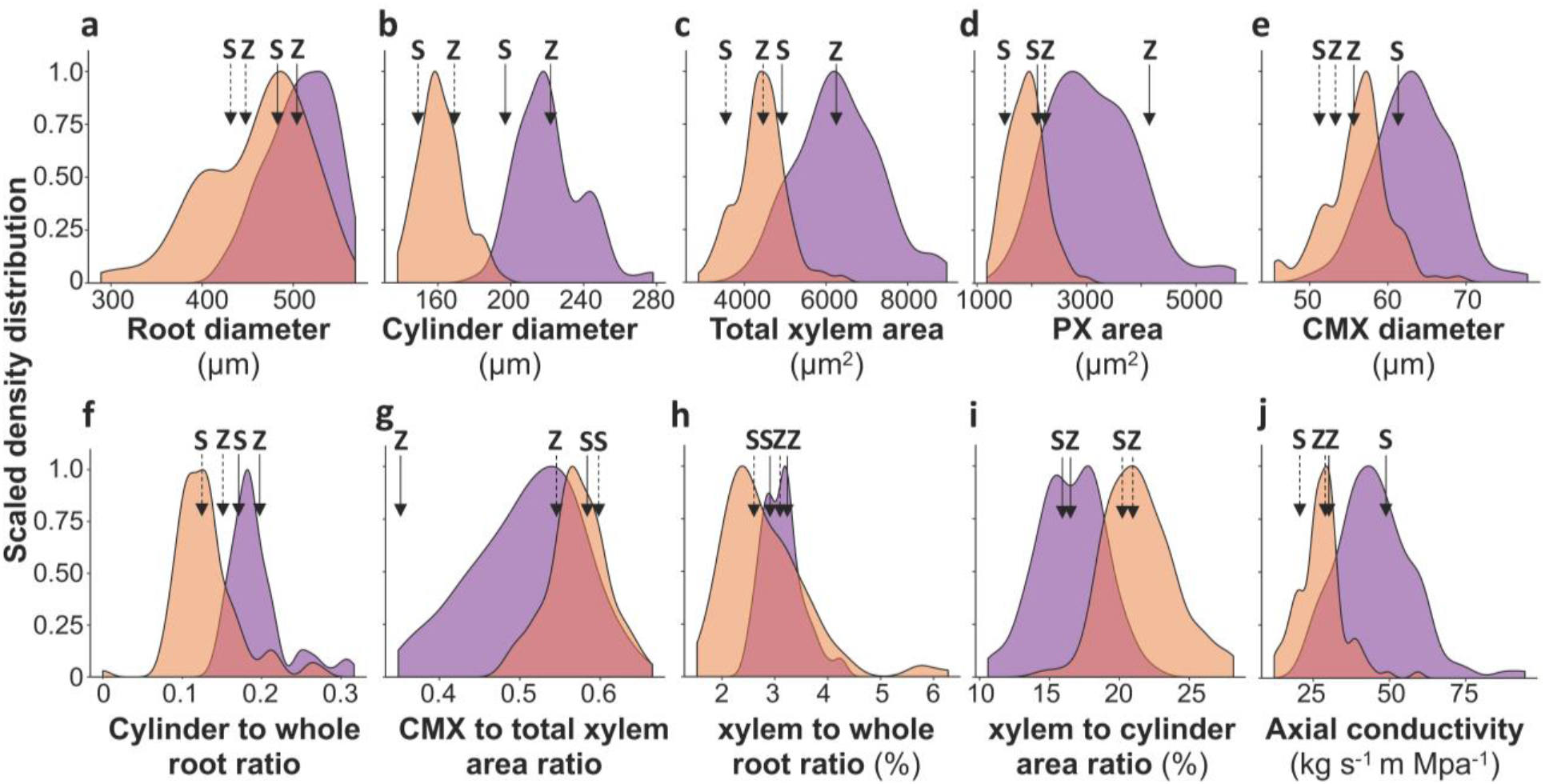
Density distribution of seminal roots anatomical and physiological traits: (**a**) root diameter (**b**) cylinder diameter, (**c**) total xylem area, (**d**) peripheral xylem (PX) area, (**e**) central metaxylem (CMX) diameter, (**f**) cylinder to whole root ratio, (**g**) CMX to total xylem area ratio, (**h**) xylem to whole root ratio, (**i**) xylem to cylinder area ratio and (**j**) axial conductance (Kx), in 150 recombinant inbreed lines (Svevo × Zavitan) for root tip (orange) and root base (violet). Mean values of the parental lines, Svevo (S) and Zavitan (Z) are indicated by dashed (root tip) and solid (root base) arrows.

### Inter-relationships between anatomical, morphological, and physiological root traits

To understand the relationships between the various seminal root morphological, anatomical, and physiological traits, and test the effect of the developmental stages between the root tip and base, we performed correlation analysis. The root morphological traits (primary root length, root hair length, and root hair density) were not associated with either root base or tip anatomical traits, neither with themselves (Fig. 5; Table S3). Correlation between anatomical and physiological traits in root base and root tip showed that while some traits maintained similar pattern [Kx (r=0.52), total xylem area (r=0.65), and xylem to cylinder area ratio (r=0.69)] others had different behavior [root diameter (r=0.16), CMX to total xylem area ratio (r=0.20), and xylem to whole root ratio (r=0.20)] (Fig. 5; Table S4). Base Kx was positively correlated with the anatomical traits in root base: CMX diameter (r=0.93, *P*=1×10^−4^), total xylem area (r=0.76, *P*=1×10^−4^) and root diameter (r=0.53, *P*=1×10^−4^). Tip Kx showed a similar pattern, except for root diameter (Fig. 5; Table. S3). While Kx is a consequence of the total xylem area (Hagen–Poiseuille’s equation), the larger elements can have a strong effect on the outcome value. Notably, CMX to total xylem area ratio showed no correlation with Kx (r=0.1 and 0.2, for root base and root tip, respectively; Table S3).

**Figure 5.**
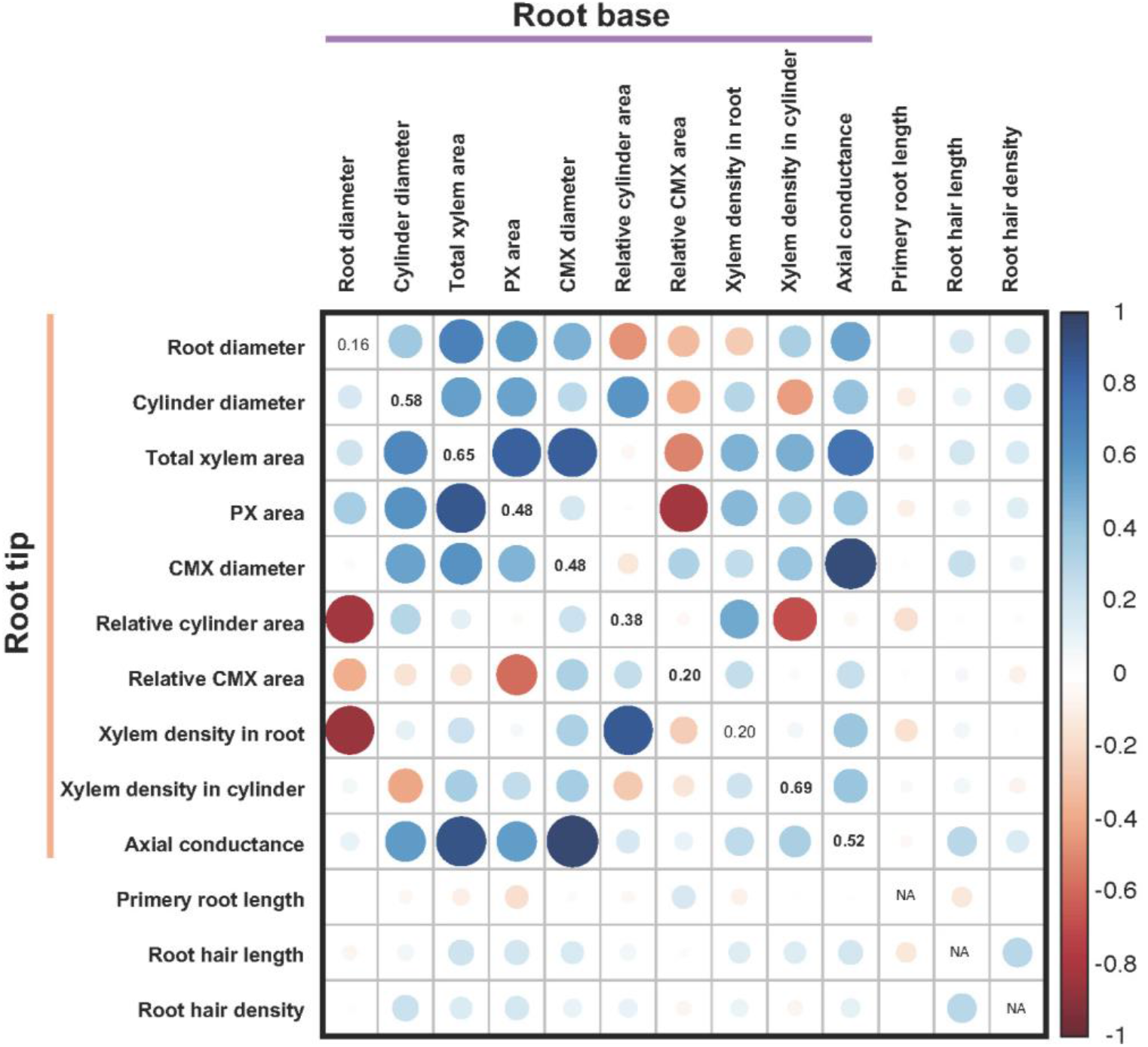
Pearson correlation matrix between the seminal roots morphological, anatomical, and physiological traits: root diameter, cylinder diameter, total xylem area, peripheral xylem (PX) area, central metaxylem (CMX) diameter, cylinder to whole root ratio, CMX to total xylem area ratio, xylem to whole root ratio, xylem to cylinder area ratio, axial conductance (Kx), primary root length, root hair length, root hair density. The upper and lower triangle represents the root base and root tip traits, respectively. The diagonal values represent the correlation between root base and root tip traits, with significant values indicated in bold (*P*≤0.05). For the primary root length, root hair length, and root hair density there is only a single observation. Colors indicate the level of correlation (r) from positive correlation (blue) to negative (red). Circle size indicates the level of significance.

### Genetic architecture of seminal root morphological, anatomical, and physiological traits

To better understand the genetic basis of the seminal roots morphological, anatomical, and physiological traits, we conducted QTL analysis, which resulted in 75 QTL (Table 1).

**Table 1.** Summary of quantitative trait loci (QTL) detected in tetraploid wheat (Svevo × Zavitan RILs population) associated with seminal roots morphological, anatomical, and physiological traits.

| Trait | #QTL | LOD<sup>a</sup> | PEV<sup>b</sup> | Svevo<sup>c</sup> | Zavitan |
| --- | --- | --- | --- | --- | --- |
| Primary root length | 8 | 6.78-10.72 | 77.6 | 5 | 3 |
| Root hair length | 5 | 4.15-5.07 | 20.8 | 4 | 1 |
| Root hair density | 2 | 3.45-3.60 | 39.4 | 1 | 1 |
| **Total** | **15** |  |  | **10** | **5** |

| Root Base |  |  |  |  |  |
| --- | --- | --- | --- | --- | --- |
| Trait | #QTL | LOD | PEV | Svevo | Zavitan |
| Root diameter | 2 | 2.71-4.45 | 24.3 | 1 | 1 |
| Cylinder diameter | 2 | 3.39-4.23 | 24.3 | 1 | 1 |
| Total xylem area | 4 | 4.37-5.75 | 24.8 | 2 | 2 |
| Peripheral xylem | 8 | 3.92-7.79 | 35.3 | 4 | 4 |
| Central metaxylem diameter | 5 | 3.12-8.50 | 67.8 | 4 | 1 |
| Relative cylinder area | 4 | 3.83-9.34 | 51.8 | 2 | 2 |
| Relative central metaxylem area | 7 | 4.60-9.91 | 58.4 | 5 | 2 |
| Xylem density in root | 2 | 2.68-3.00 | 68.0 | 1 | 1 |
| Xylem density in cylinder | 2 | 3.01-4.07 | 23.8 | 2 | 0 |
| Axial conductance | 4 | 6.11-6.13 | 19.7 | 2 | 2 |
| **Total** | **40** |  |  | **24** | **16** |

<sup>a</sup>LOD scores that were found significant when comparing hypotheses $H_1$ (there is a QTL in the chromosome) vs. $H_0$ (no effect of the chromosome on the trait), using 1000-permutation test (Churchill & Doerge, 1994).
<sup>b</sup>Proportion of explained variance of the trait.
<sup>c</sup>The allele contributing to a higher value of each trait.
n.d., not detected.

For each trait, we detected between one QTL (tip root diameter and tip CMX to total xylem area ratio) to 8 QTL (primary root length and base PX area), with two traits (tip xylem to cylinder area ratio and tip Kx) without detected QTL. LOD score ranged from 2.68 (base xylem to whole root ratio) to 10.72 (primary root length), explaining from 14% (tip CMX to total xylem area ratio) to 77.6% (primary root length) of the phenotypic variance. For all QTL we considered the higher value as additive. In general, for most QTL, the Svevo allele conferred higher values (42 *vs*. 33), with 41 QTL located on the A sub-genome and 34 on the B sub-genome (Tables 1 and S5; Fig. 6). Genetic correlations between traits resulted in several significant associations: base root diameter and tip PX area (*P*=0.01), base xylem to cylinder area ratio with tip PX area (*P*=0.01), tip cylinder to whole root ratio, and tip xylem to whole root ratio (*P*<10^−4^), and primary root length and base CMX diameter (significance at *P<*0.01) (Table S6).

**Figure 6.**
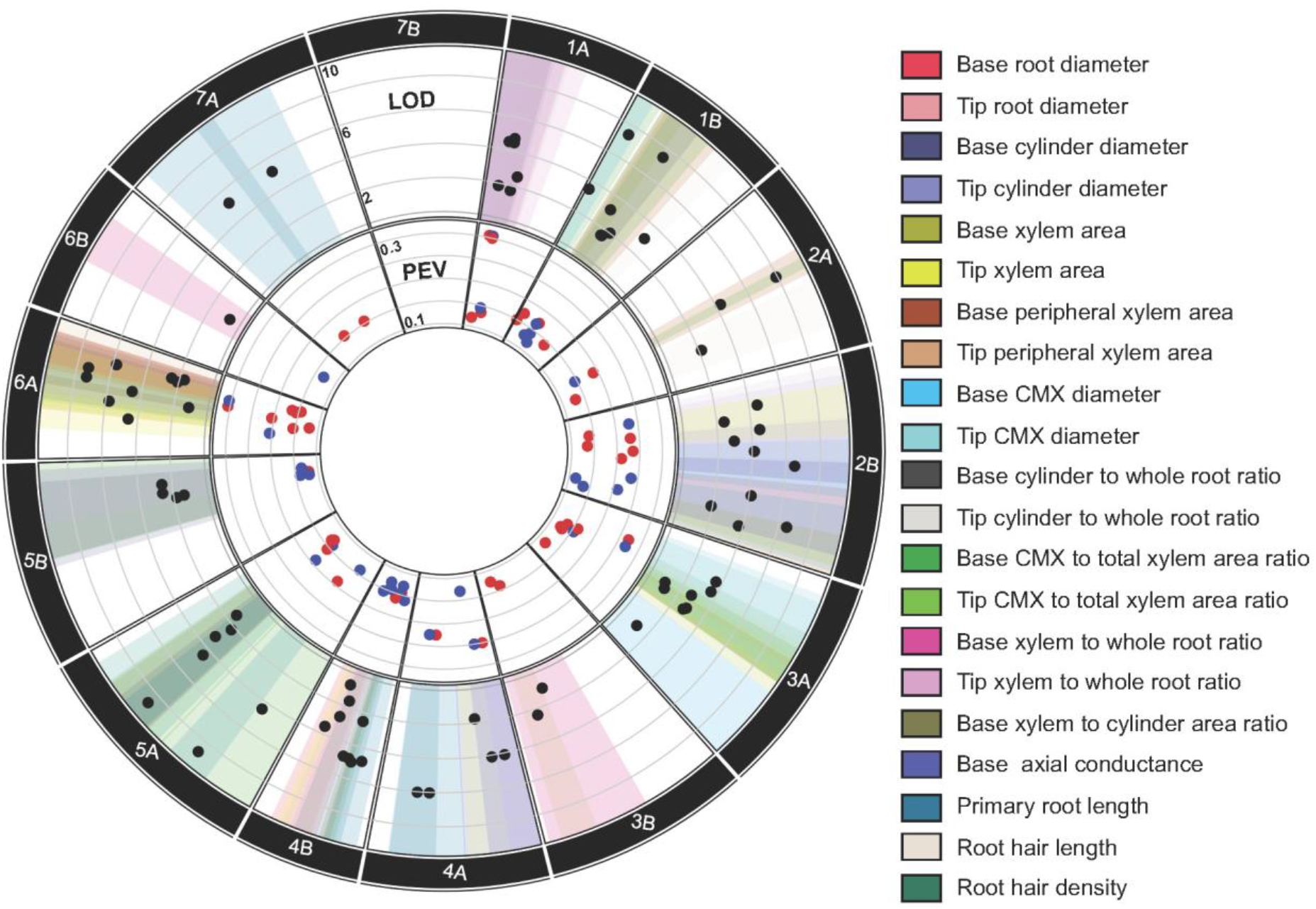
Quantitative trait loci (QTL) associated with 21 seminal root traits in recombinant inbred lines (RILs) of the cross between Svevo and Zavitan. Traits were analyzed in the root tip and based for *morphological*: primary root length, root hair length, and root hair density; *anatomical*: root diameter, cylinder diameter, total xylem area, peripheral xylem (PX) area, central metaxylem (CMX) diameter, cylinder to whole root ratio, CMX to total xylem area ratio, Xylem to whole root ratio and xylem to cylinder area ratio. *Physiological*: axial conductance (Kx). Circos diagram showing, from the outer to inner tracks, the wheat chromosomes, QTL for specific traits and their positions, LOD score (black markers), percentage of explained variance (PEV) where Svevo (grey) and Zavitan (red) additive contribution are marked.

Major genomic regions associated with several overlapping QTL, i.e. hotspots, can indicate either cluster of several genes with a different function or a major gene with a pleiotropic effect. We detected two hotspots with overlapping QTL (additive effect marked for the higher allele, Svevo and Zavitan). On chromosome 5A (564,800,028-672,857,031 bp), primary root length (Z), root hair length (S), base CMX to total xylem area ratio (Z), base PX area (S), base cylinder diameter (S). On chromosome 6A (564,840,388-614,128,384 bp) consist five QTL: base root diameter (S), base cylinder to whole root ratio (Z), base xylem to cylinder area ratio (S), base PX area (S), and tip PX area (S) (Fig. S5)

### Validation of ancestral axial conductance related alleles

Zavitan accession was collected from its natural habitat with shallow brown basaltic soil type, which has high soil moisture fluctuation (Peleg et al., 2008). Thus, we targeted wild alleles that could be important for reshaping the root architecture to cope with fluctuating soil moisture for further analysis. To validate the major root base QTL affecting Kx, and CMX diameter, we used wild emmer wheat Zavitan introgression lines in the background of elite durum wheat Svevo. Two QTL conferring base Kx on chromosomes 2B (Svevo) and 4A (Zavitan) were tested using IL with opposite genetic composition: IL108 (Zavitan and Svevo) and IL95 (Svevo and Zavitan, respectively). Overall, our results validated the QTL composition, with IL95 exhibiting significantly higher base Kx (54.75 *vs*. 44.49) compared with Svevo, whereas IL108 had a significantly lower base Kx (34.77) (Fig. 7).

**Figure 7.**
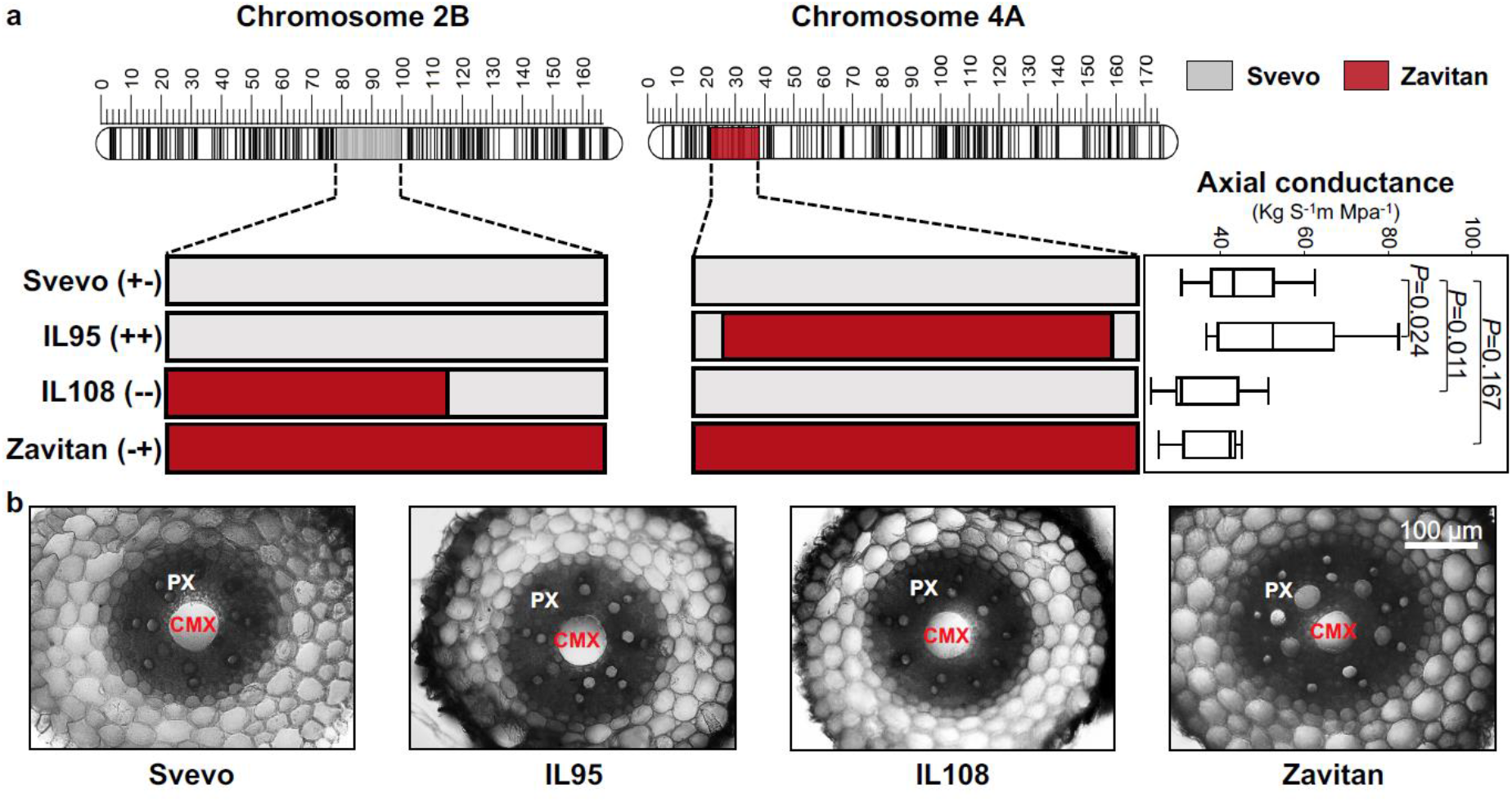
Validation of QTL for seminal root axial conductance (Kx). (**a**) Graphical genotyping of the two parental lines (Svevo and Zavitan) and two wild emmer introgression lines (IL108 and IL95) harboring Kx QTL on chromosomes 2B and 4A, respectively. The Box plot represents each genotype Kx and comparison to the recurrent parent Svevo according to Dunnett test (*n*=8). (**b**) A represented image of seminal root base anatomical cross-section of each genotype. The central metaxylem (CMX) and peripheral xylem (PX) are marked.

We validated two base CMX diameter QTL (on chromosomes 1B and 6A), where Svevo contributed the adaptive allele, using IL100 and IL64 that harbor the Zavitan allele, respectively. Both ILs as well as Zavitan, exhibited significantly lower (∼12%) CMX diameter compared to Svevo (Fig. 7). Notably, while CMX diameter has a pivotal role in determining Kx (as expressed in Hagen-Poiseuille’s equation), most CMX diameter QTL did not overlap Kx QTL.

## DISCUSSION

Enhancing tolerance to water stress through breeding is essential for maintaining production under the projected climate change (Anderson, Bayer, & Edwards, 2020). While genetic dissection of various above-ground traits has greatly improved our knowledge (Salvi & Tuberosa, 2015), in recent years more emphasis has been given to the hidden half of the belowground root system. Most of these studies focus on the root system architecture morphology [i.e. root length, density, and angle (Lynch, 2019; Voss-Fels et al., 2018; Uga, 2021)], aiming to discover QTL with the potential to improve water and nutrient acquisition under suboptimal environments. However, the genetic basis of root anatomical and physiological traits were less studied (Schneider & Lynch, 2020).

### Form lab-to field-based high-throughput root traits phenotyping

The “out-of-sight” nature and extreme phenotypic plasticity of roots under native field conditions (due to dynamic responses to varying water, nutrient, and environmental cues) and the limited availability of high-throughput phenotyping tools, inhibited functional genetic studies of root architecture and especially anatomical and physiological traits (Atkinson, Pound, Bennett, & Wells, 2019; Watt et al., 2020). Moreover, the opaque nature of the soil further increases the complexity of *in-situ* phenotyping of root systems. While controlled (i.e. hydroponic) root phenotyping was not always a good predictor of the field situation (Rich et al., 2020), Alahmad et al. (2019) showed that root angle phenotyping in the lab was consistent with the expression in the field. Likewise, seminal root number phenotypes observed in hydroponic conditions are transferrable to soil conditions, especially during the vegetative phase (Golan et al., 2018; Richard et al., 2015; Watt et al., 2013).

For this study, an important question was to determine if the young seedling assay for phenotypic differences can be observed for a longer duration in the field. Our wide screening of young seedlings using the cigar-roll method (i.e. on germination paper) in control conditions enabled us to identify genotypes with contrasting Kx (Svevo and IL82). Further field-based analysis revealed that although the absolute values were larger compared with the lab experiment, a similar trend of ∼50% difference between the genotypes was maintained throughout development (Figs. 2, S3, S4). Thus, the phenotypic approach developed for the current study can represent the field performance in terms of axial conductance and related anatomical traits.

### Root morphological traits are independent of other root properties

Under the semi-arid Mediterranean climate, soil moisture in the upper layer is limited and young seedlings are exposed to frequent events of dehydration. Recently, we showed that during wheat evolution under domestication, the number of seminal roots increased (from 3 to 5) due to the activation of two root meristems (Golan et al., 2018). While one can expect that the difference in the number of seminal roots will affect morphological and anatomical root traits, surprisingly, we could not find any QTL overlapping with this trait. The fast development of seminal roots is important for reaching deeper soil layers to avoid the dehydration associated with the erratic rain distribution of the Mediterranean basin. Eight QTL associated with primary root (SR1; Fig. 1) length were found in the current study (Table S5), explaining most (76%) of the phenotypic variance. Since both parental lines have full genome sequence, we were able to test genetic overlaps with previous studies and found three genomic regions on chromosomes 4A, 4B, and 7A (Maccaferri et al., 2016), whereas the other five QTL (chromosomes 4A, 4B, 5A, 5B, and 7A) are novel.

The proliferation of root hairs along major root axes has been implicated in a range of processes including, enhanced water uptake and nutrient acquisition, root-microbial signal exchange, allelochemical release and plant anchoring (Carminati et al., 2017; Holz, Zarebanadkouki, Kuzyakov, Pausch, & Carminati, 2018). While all these processes are associated with the contribution of root hair to extending the effective radius surface area of the root, existing results on their function are not fully understood. Moreover, quantification of root hairs features is challenging as the trait is highly affected by environmental conditions (Salazar-Henao, Vélez-Bermúdez, & Schmidt, 2016). As consequence, the genetic basis of root hairs was less studied, and the limited available research tested only root hair length (Horn, Wingen, Snape, & Dolan, 2016; Liu et al., 2017).

Here we dissect root hair characteristics into length and density in an attempt to represent the complexity of the tissue. The different contribution of root hair length and density is exemplified in the comparison between the two parental lines of our mapping population. While the wild parental (Zavitan) had short root hairs compared to the domesticated parent (Svevo), its density was much higher (Fig. 3b, c). In general, our genetic analysis resulted in the identification of five QTL for root hair length and two QTL for root hair density (Fig. 6; Table. S5). Genetic analysis of root hair length in hexaploid populations [bread wheat (Horn et al., 2016) and spelt wheat (Okano et al., 2020)] resulted in 4 and 1 QTL, respectively. Only one of these QTL (on chromosome 2A) co-localized with the five root hair length QTL found in the current study for tetraploid wheat, which further emphasizes the potential of finding new alleles in wild wheat germplasm. Genetic characterization of root hair density in wheat or other grasses remains unexplored. It is worth noting that in contrast with our initial hypothesis, low correlation was found between root hair traits and other anatomical traits (r<0.3; Fig. 5). Thus, it is yet to be discovered how root hair length and density interact with other root trait and their collective effect on water uptake.

### Genetic architecture of xylem elements suggests independent control among young and mature root tissue

Xylem tracheary elements have a unique life cycle; from cell differentiation in the root tip to rapidly programmed cell death as they reach their full functional state (Fukuda, 1997), which makes it difficult to track precisely each developmental stage. To study the root anatomical traits, we analyzed the root tip, which represents the differentiating stage, and the root base, which represents mature cells. In accordance, we found that cylinder diameter and xylem elements increased from tip tissue to the root base as indicated by the significant positive correlation between tissues (Figs. 4b-e, 5). Generally, we were able to detect more QTL for anatomical traits in the root base compared with tip, and interestingly most QTL did not co-localize, which suggests distinct genetic regulation of xylem development on a spatial scale (Fig. 6; Table. S5).

In temperate cereal roots, axial conductance (Kx) was found to be low in the tip differentiation zone and steady along the mature root tissue (Bramley, Turner, Turner, & Tyerman, 2009; Knipfer & Fricke, 2010). In our analysis, we were able to detect QTL for CMX and peripheral xylem at the tip, which are the components of axial conductance, however, no root tip Kx QTL was detected (Table 1; Fig. 6). The absence of tip Kx QTL detection may be a consequence of the fact that xylem elements reach maturity after 96 h (Fukuda, 1997). Our results support Fukuda’s findings where phenotyping these developing xylem elements result in a strong environmental effect as expressed in a higher coefficient of variance (CV=27, CV=7.2, and CV=17.6 for Kx, CMX, and PX, respectively). Nevertheless, our physiological measurements indicated that Kx in root tip was strongly correlated with base Kx (r=0.52, *P*<0.0001; Table S4), therefore we decided to focus on the root base Kx as the representative of root potential axial conductance. It is worth noting that while axial conductance values calculated from the vessel’s diameters can differ from the actual Kx (Tyree & Ewers, 1991), they were found to correlate to each other. (Strock, Burridge, Niemiec, Brown, & Lynch, 2020; Martre, Durand & Cochard, 2000). Therefore, the values obtained from the cross-sections can credibly represent the actual hydraulic differentness between genotypes, especially when comparing within the same species. Furthermore, several studies have shown an association between the xylem vessel dimensions and physiological performance of the whole plant (Tombesi, Johnson, Day & Dejong, 2010; Schoppach, Wauthelet, Jeanguenin & Sadok, 2014; Bheemanahalli, Hechanova, Kshirod & Krishna Jagadish, 2019).

### Reshaping root hydraulics via axial conductance QTL

Lower root Kx was identified as a key trait supporting sustainable grain yield under terminal drought, subsequent of reduced water uptake during the early stages of growth (Richards & Passioura, 1989). Likewise, our field trial resulted in a 9% yield advantage for IL82 with lower Kx (30%) under terminal drought (Fig. 2; Table S2). In contrast to Richards and Passioura’s (1989) findings, our results suggest that IL82 had lower yield under well water conditions likely due to a yield penalty associated with lower Kx (Table S2). Although Passioura’s concept was introduced ∼40 years ago (Passioura, 1983), and repeatedly discussed in the literature (e.g. Berger, Palta, & Vadez, 2016; Blum, 2009; Richards, Rebetzke, Condon, & van Herwaarden, 2002), the genetic basis of Kx was not studied yet. Here we integrated a new high-throughput root anatomical cross-section pipeline with a population derived from a cross between wild and domesticated tetraploid wheat to determine the genetic basis of root Kx. Four novel genomic regions associate with root Kx were identified, which explain most of the phenotypic variance (62%). A meta-analysis of these genomic regions exposed co-localization with QTL for grain yield on chromosome 2B in a population of durum wheat × wild emmer, under water-limited conditions (Peleg et al., 2009) (Table S7). Notably, both alleles are derived from the wild emmer accessions (Zavitan reduce Kx and G18-16 increased yield, respectively).

Wild emmer has evolved over long evolutionary history under the fluctuating precipitation of the Mediterranean basin and accumulated ample genetic diversity for drought adaptations (e.g. Golan et al., 2018; Peleg et al., 2009). Under such conditions, narrow xylem elements can contribute to the natural population survival and seed set. Introgression of wild Kx allele (Zavitan) into Svevo background further validates the potential of wild alleles for redesigning seminal root anatomy and hydraulic properties of modern wheat with increased resilience (Fig. 7). However, root hydraulics is a complex trait associated not only with Kx but also with radial conductivity and root architectural traits that were not examined in the current study. Since roots are highly plastic, it remains to be determined, how altering one specific allele could impact the compensating response among other traits.

### Xylem elements spatial distribution affect its hydraulic properties

Under water-limiting environments, xylem vessel diameter and distribution affect plant resistance to embolisms (Lucas et al., 2013). Anatomical characterization of the two parental lines revealed a distinct xylem vessel distribution in Zavitan, with narrow CMX and larger PX (Fig. 7b). Prompted by this observation, we integrated a new index of CMX to total xylem area ratio. Notably, Zavitan expressed the lowest values compared to the whole RILs population (Fig. 4g). Genotypes which have similar peripheral xylem area differ dramatically in the CMX diameter and *vis versa* (Fig. S7). Moreover, most QTL associated with CMX and PX did not show genomic co-localization, which suggests that there is a possibility to break the compensating association between both traits.

From the hydraulic perspective, this ratio between CMX and PX could also play a role in reducing the risk for cavitation under drought. Cavitation occurs when the tension in the vessels increases to a point of air bubbles formation and water column break. This phenomenon may occur due to exposure to periods of severe drought and can result in loss of function of the whole vessel. It is suggested that wider vessels are at higher risk of cavitation (Melvin, Stephen, & Hervè, 1994; Strock, Burridge, Niemiec, Brown, & Lynch, 2020), hence plants with lower CMX area may be less susceptible to cavitation. Moreover, even in the case of CMX failure, plants with larger PX (as found in the wild accession Zavitan) have the potential to meet transpiration demand. It is worth noting that IL82, which had a higher grain yield under terminal drought and low Kx, also has a significantly lower CMX to total xylem area ratio compared with Svevo (*P*=0.002). Collectively, our results suggest that the CMX to total xylem area ratio should be considered a target trait for breeding drought resilience cultivars.

## Conclusion and future perspective

Here we tested the hypothesis that under the terminal drought conditions associated with the Mediterranean basin, smaller xylem vessels decrease water-use, thus supporting more residual water for the critical phase of grain filling (Passioura, 1972). A field-based characterization further supported the notion that lower Kx promotes sustainable grain production under terminal drought conditions. We harnessed the wide genetic diversity of wild emmer wheat to uncover the genetic basis of Kx and associated morphological, anatomical, and physiological seminal root traits. To the best of our knowledge, this is the first report on QTL associated with seminal roots anatomical and physiological traits in wheat. Our results shed new light on the structural and functional anatomy of seminal roots and their association with yield sustainability under suboptimal environments. Integration of the identified wild alleles into future breeding programs may facilitate the optimization of seminal root hydraulic properties and should be considered while shaping the development of new wheat (and other cereals) cultivars for the projected climate change environments.

## Acknowledgment

We thank members of the Peleg lab for helpful comments during the preparation of this work. We would like to thank N. Teboul for her assistance with the fine mapping, Dr. R. Hayouka for her technical support with the phenotyping experiments, and L. Shemesh for drawing the seminal roots illustration. This study was partially supported by the Israel Ministry of Agriculture and Rural Development (grants # 20-10-0066; 12-01-0005), and the U.S. Agency for International Development Middle East Research and Cooperation (grant # M34-037).

