## Supplementary material for "Deciphering the genetic basis of wheat seminal root anatomy uncovers ancestral axial conductance alleles": SI Figures

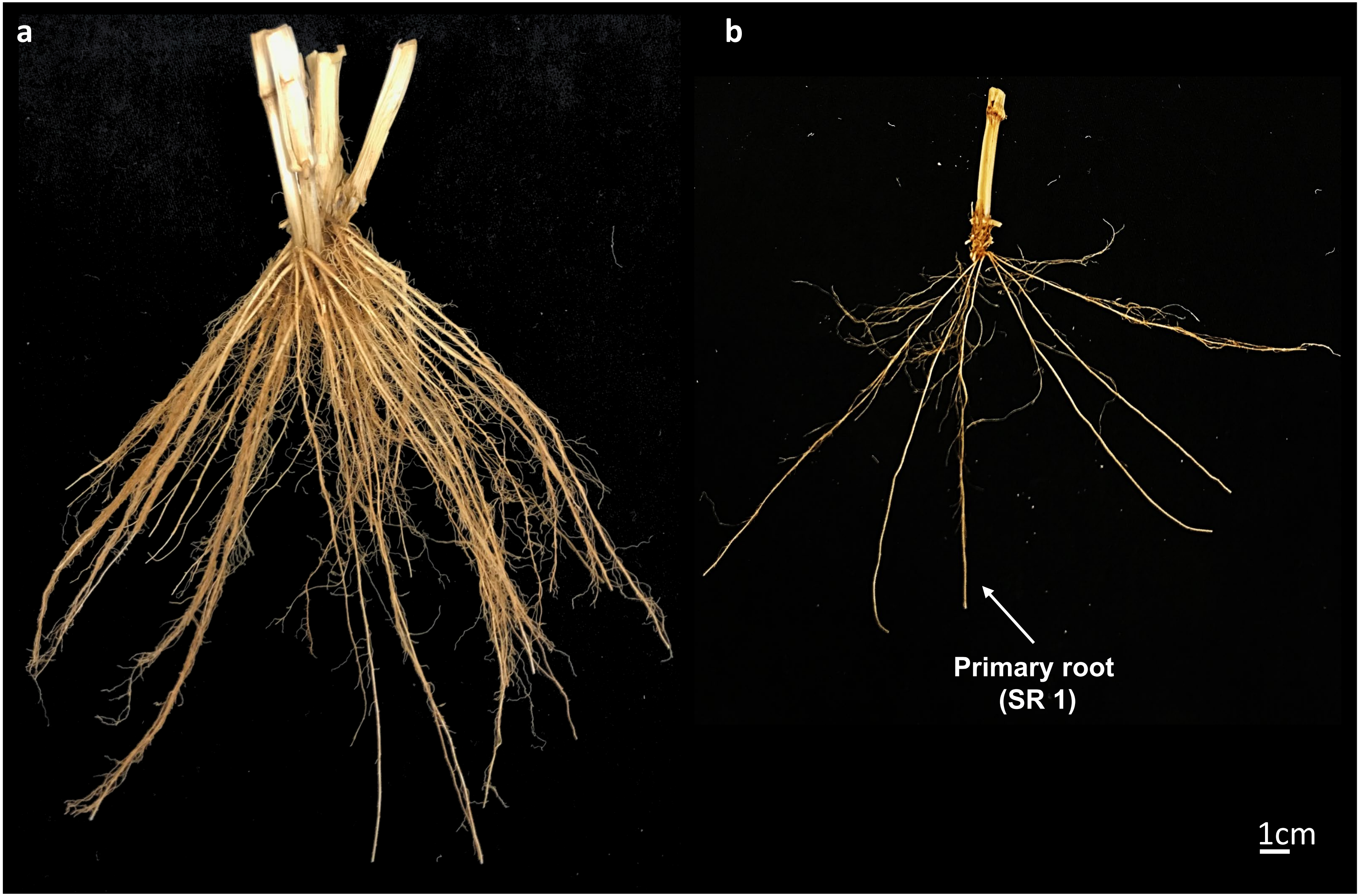


**Figure S1.** A representative plant from field experiment after harvesting. (**a**) Whole root system (at harvest) and (**b**) manual removed adventive roots to expose the seminal roots. Primary root (seminal root #1) is indicated by arrow.

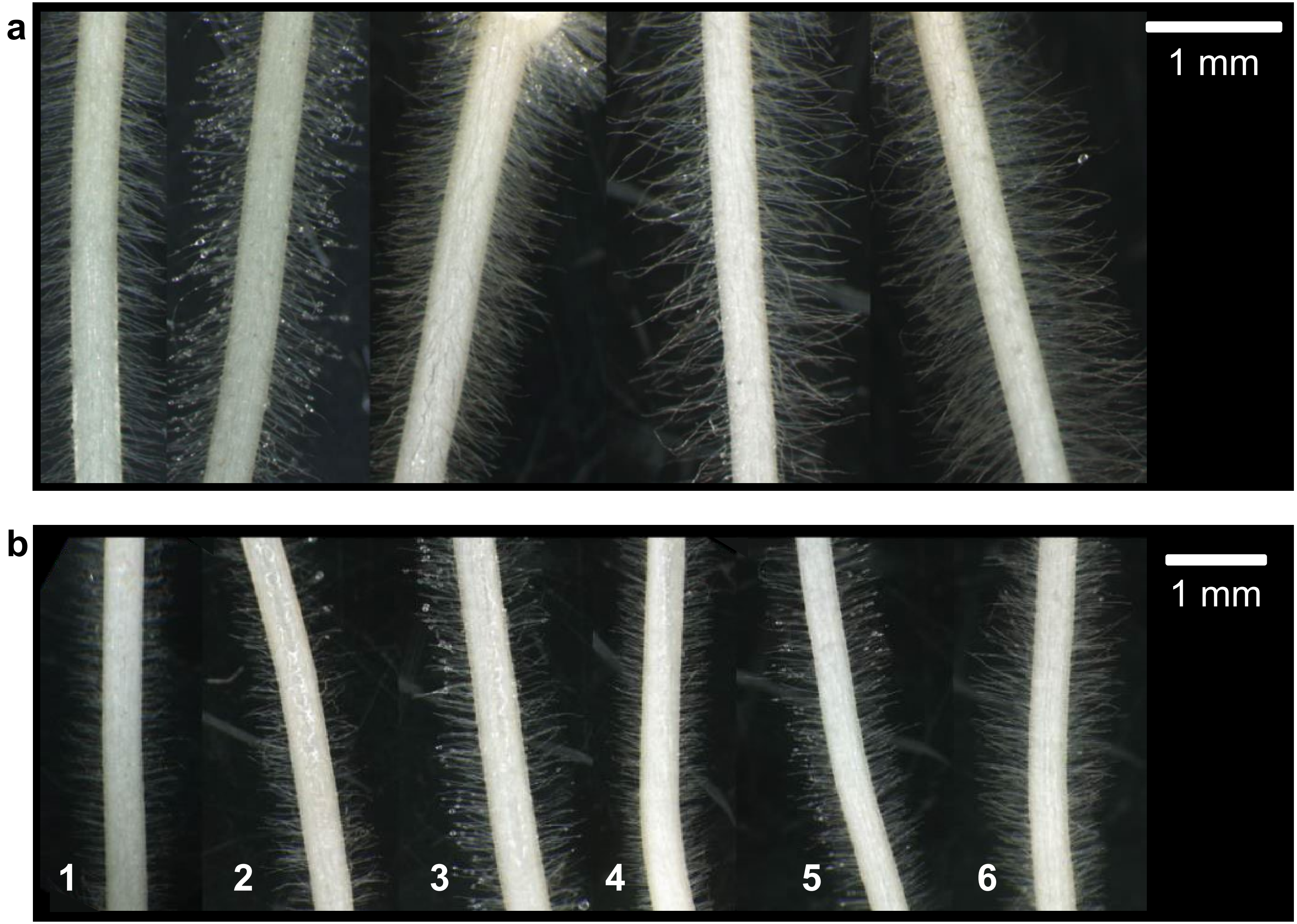


**Figure S2**. Phenotypic diversity of root hairs length and density in recombinant inbred lines (Svevo x Zavitan). (**a**) A representative panel of root hair length ranging from 330 μm (left) to 875 μm (right). (**b**) Root hairs density index range from 1 (low) to 6 (high).

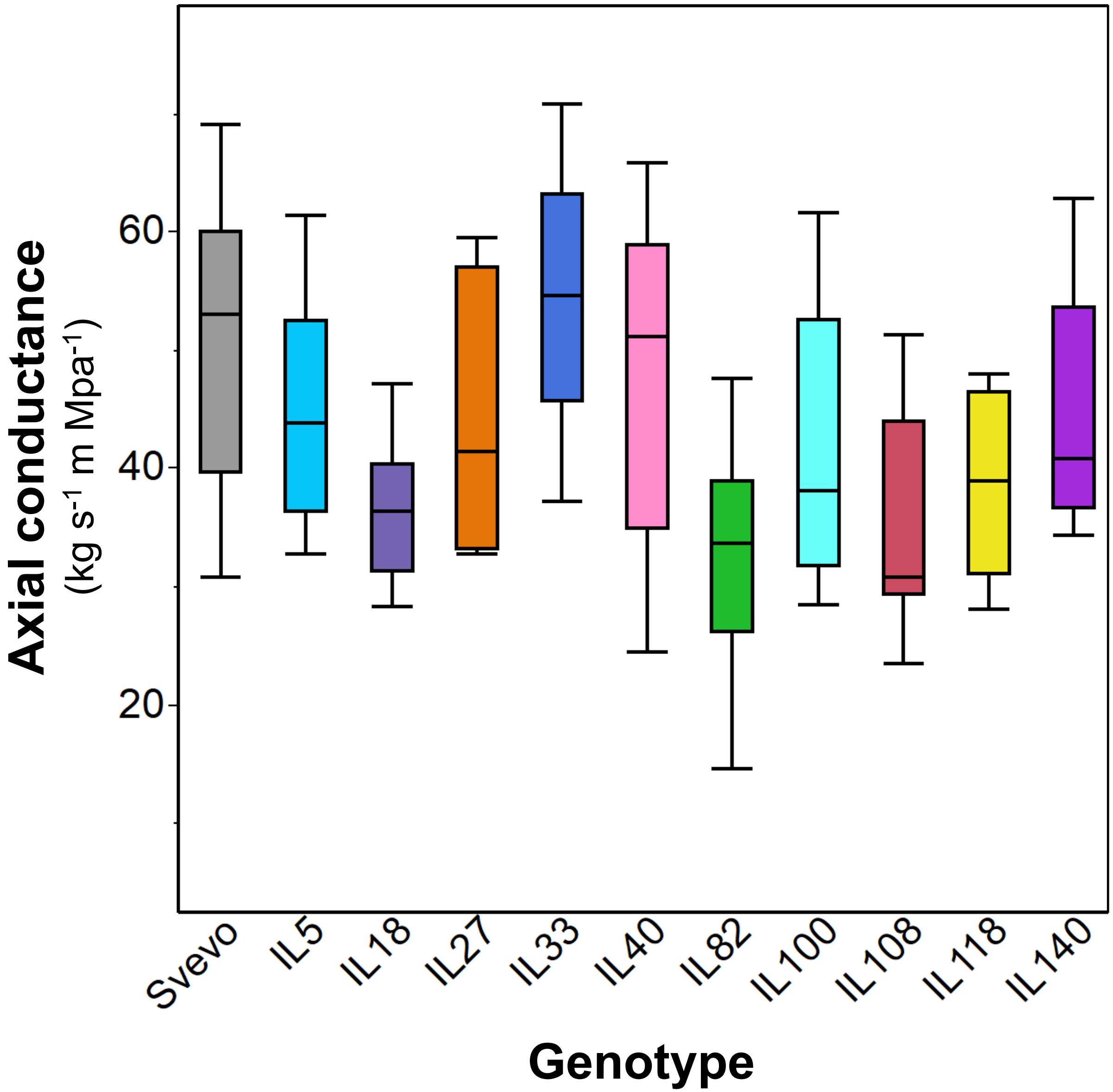


**Figure S3.** Seminal root base axial conductance of 10 wild emmer introgression lines (ILs) and their recurrent parent Svevo under control conditions (*n*=7).

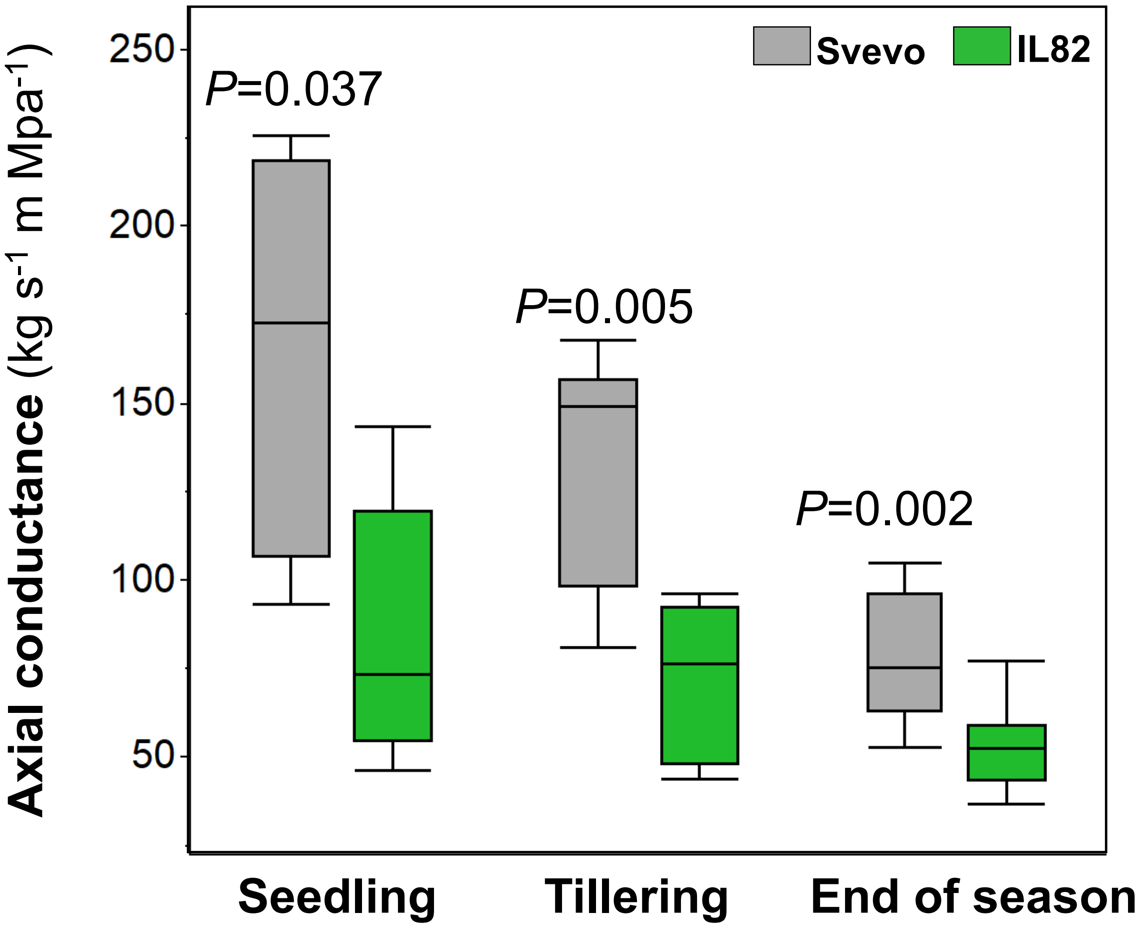


**Figure S4.** Root base axial conductance of Svevo and IL82. Measurements were taken at three developmental stages: seedling (10 days after sowing, DAS), tillering (34 DAS), and end of the growing season (140 DAS). Differences between genotypes were analyzed by t-test ($n$=5).

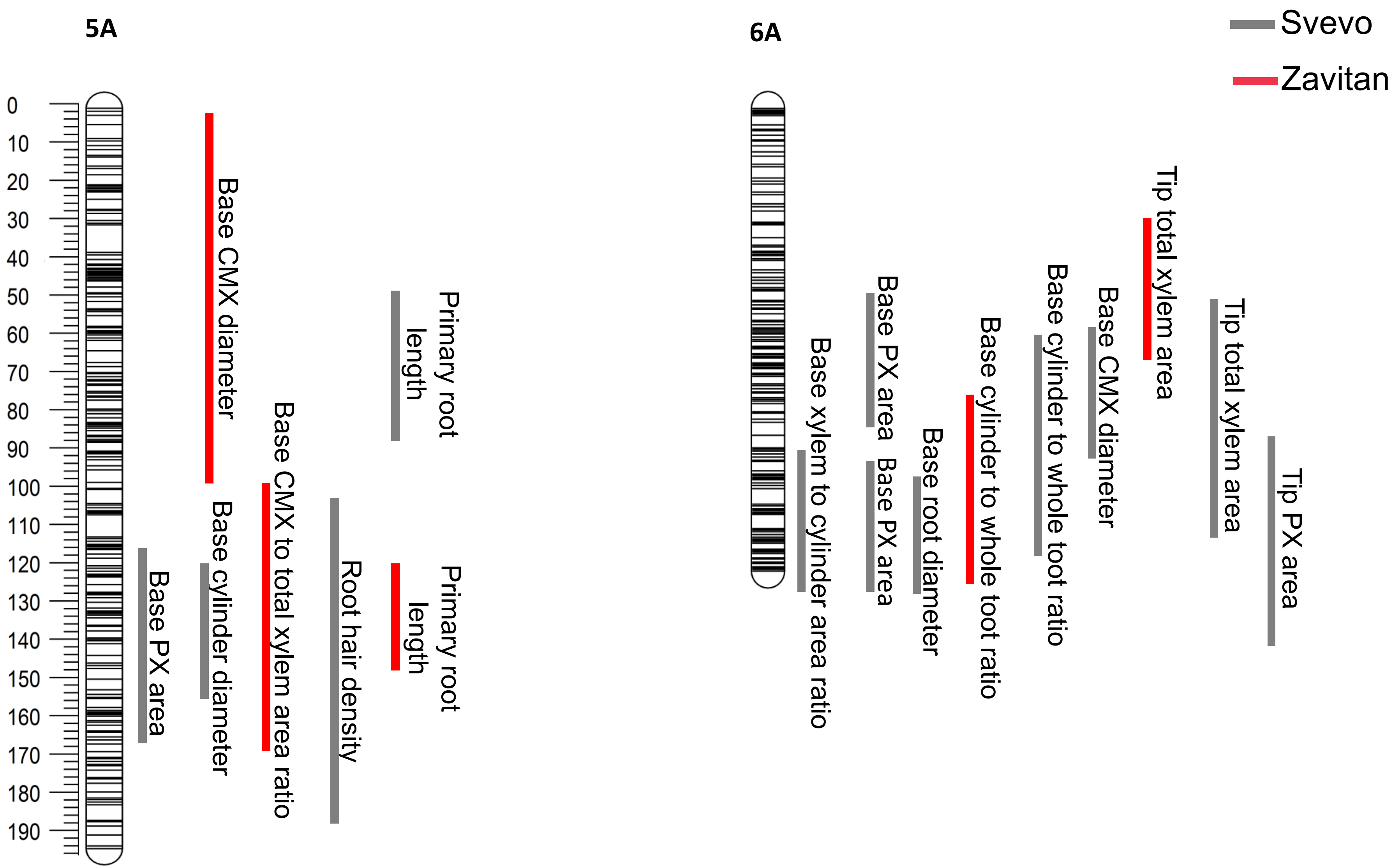


**Figure S5.** Co-localization of QTL on chromosomes 5A and 6A. Traits were analyzed in the root tip and based for: primary root length, root hair density, root diameter, cylinder diameter, total xylem area, peripheral xylem (PX) area, central metaxylem (CMX) diameter, cylinder to whole toot ratio, xylem to cylinder area ratio, and CMX to total xylem area ratio. QTL color represents the allele with a higher value.

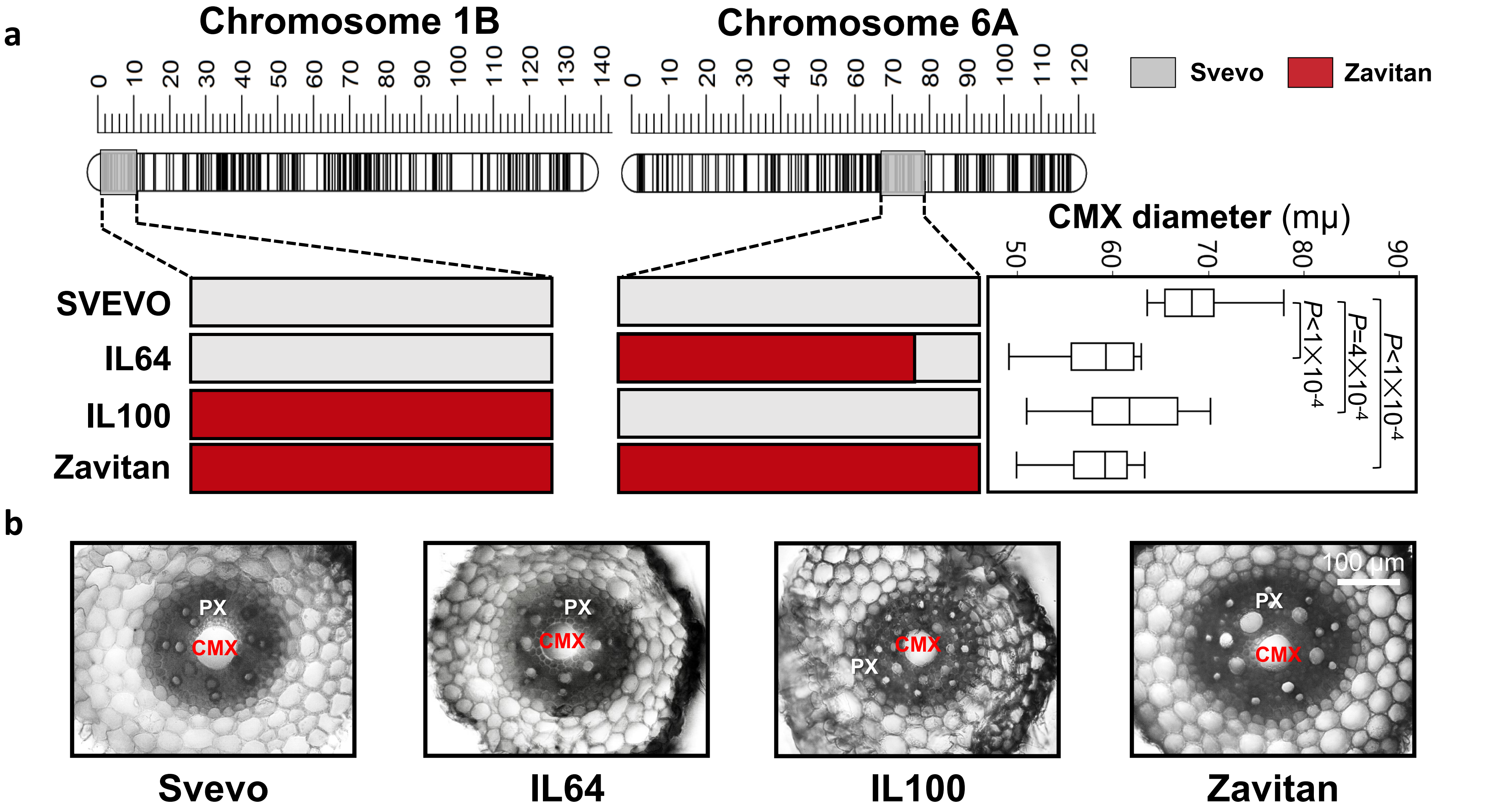


**Figure S6.** Validation of QTL for seminal root central metaxylem (CMX) diameter. (**a**) Graphical genotyping of the two parental lines (Svevo and Zavitan) and two wild emmer introgression lines (IL100 and IL64) harboring CMX diameter QTL on chromosomes 1B and 6A, respectively. Box plot represents each genotype CMX diameter in comparison to the recurrent parent Svevo according to Dunnett test (*n*=8). (**b**) A represented image of seminal root base anatomical cross-section of each genotype. The central metaxylem (CMX) and peripheral xylem (PX) are marked.

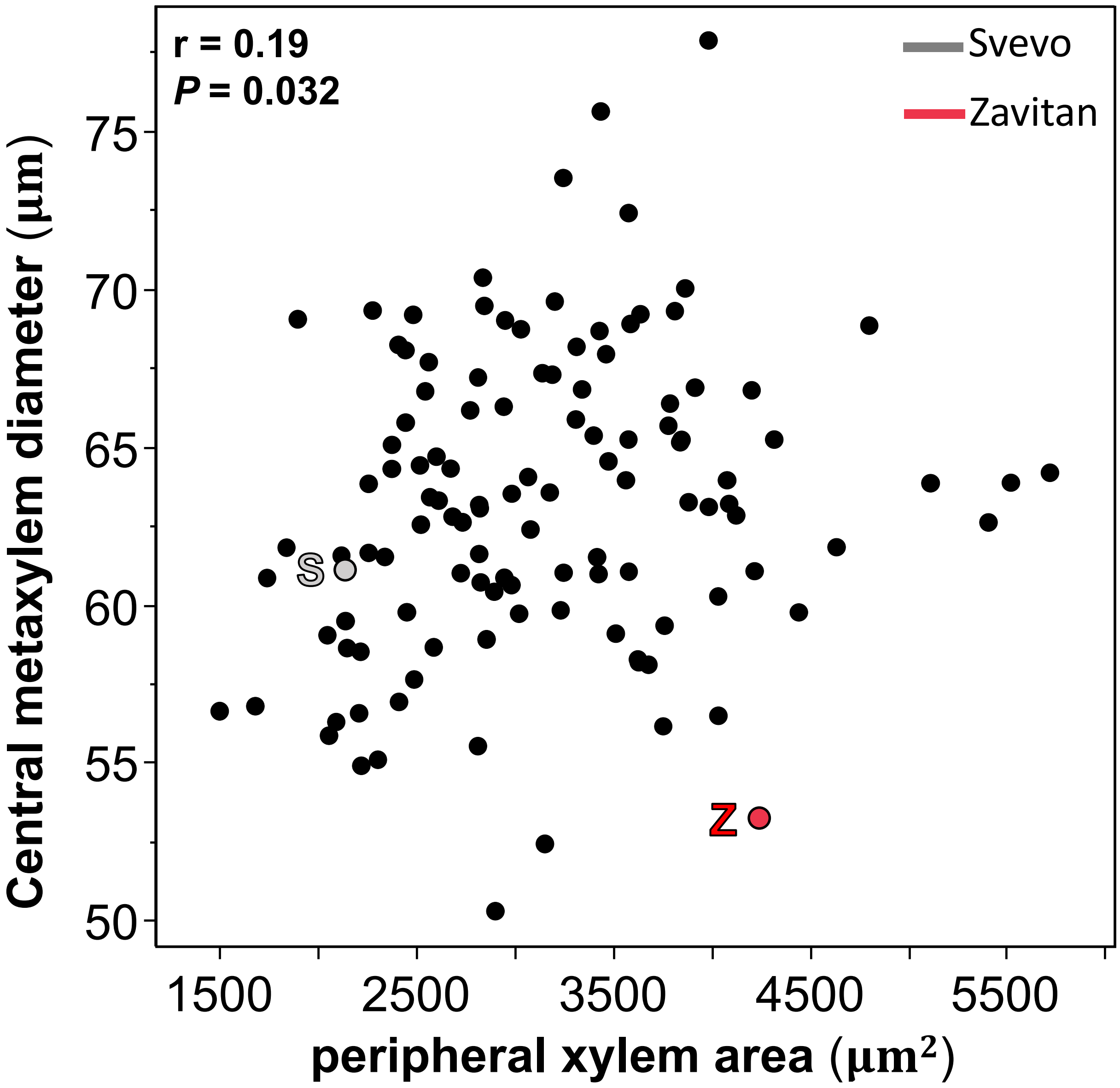


**Figure S7.** Correlation between seminal root base central metaxylem diameter and peripheral xylem area. Data is mean (*n*=4). The two parental lines, Svevo and Zavitan are marked.
